# The effects of acute Methylene Blue administration on cerebral blood flow and metabolism in humans and rats

**DOI:** 10.1101/2022.09.10.507418

**Authors:** Nisha Singh, Eilidh MacNicol, Ottavia DiPasquale, Karen Randall, David Lythgoe, Ndabezinhle Mazibuko, Camilla Simmons, Pierluigi Selvaggi, Stephanie Stephenson, Federico E Turkheimer, Diana Cash, Fernando Zelaya, Alessandro Colasanti

## Abstract

Methylene Blue (MB) is a brain-penetrating drug with putative neuroprotective, antioxidant and metabolic enhancing effects. *In vitro* studies suggest that MB enhances mitochondrial complexes activity. However, no study has directly assessed the haemodynamic and metabolic effects of MB in the human brain.

We used *in vivo* neuroimaging to measure the effect of MB on cerebral blood flow (CBF) and brain metabolism in humans and in rats. Two doses of MB (0.5 and 1 mg/kg in humans; 2 and 4 mg/kg in rats; iv) induced reductions in global cerebral blood flow (CBF) in humans (F_(1.74, 12.17)_5.82, p=0.02) and rats (F_(1,5)_26.04, p=0.0038). Human cerebral metabolic rate of oxygen (CMRO_2_) was also significantly reduced (F_(1.26, 8.84)_8.01, p=0.016), as was the rat cerebral metabolic rate of glucose (CMRglu) (t=2.6_(16)_ p=0.018).

This was contrary to our hypothesis that MB will increase CBF and energy metrics. Nevertheless, our results were reproducible across species and dose dependent. One possible explanation is that the concentrations used, although clinically relevant, reflect MB’s hormetic effects, i.e., higher concentrations produce inhibitory rather than augmentation effects on metabolism. Additionally, here we used healthy volunteers and healthy rats with normal cerebral metabolism where MB’s ability to enhance cerebral metabolism might be limited.

## Introduction

Alterations in mitochondria-dependent energy production have been implicated in the pathophysiology of numerous neurological and psychiatric diseases, including Alzheimer’s and Parkinson’s disease, multiple sclerosis, bipolar disorder and schizophrenia ^1–6^. Compounds used as routine clinical treatments for these conditions do not address the associated neuro-energetic deficits or reverse the observed metabolic decline. This unmet need led to a recent growth of interest in identification of neuroprotective strategies aiming to revert the dysfunction of neuronal mitochondrial respiration ^7–10^. To this end, one plausible strategy is to enhance cerebral oxygen consumption via boosting the mitochondrial oxidative phosphorylation and adenosine triphosphate (ATP) production.

Methylthioninium chloride, also known as methylene blue (MB), is highly brain penetrant ^11, 12^ and hypothesised to have brain metabolic enhancing and neuroprotective properties. MB is one of the oldest known drugs, first synthesised in 1876, as a blue chemical dye and since used as a medical dye for pre-surgical staining ^13^. Other uses for MB ^11^ include that as antimalarial agent, treatment for methemoglobinemia, cyanide poisoning ^14^ and vasoplegic syndrome ^15^, as well as a more novel indication as tau-aggregation inhibitor, for treatment of Alzheimer’s Disease, that has been extended to clinical trials ^16^. Converging evidence from newer human and animal research studies suggest that MB also has antidepressant, anxiolytic, antipsychotic and mood-stabilising properties ^17^. In accordance with this proposed ‘neurotropic’ efficacy, MB was also shown to dose-dependently enhance cognitive performance, learning and memory in both humans and in experimental animals ^18–28^.

It is likely that many of MB clinical properties rest on its ability to act as an alternate electron acceptor in the metabolic electron transport chain, increasing mitochondrial oxygen consumption and boosting ATP production ^29^. These effects of MB are dose-dependent, yet not linear: lower doses (0.5-4 mg/kg, calculated based on animal studies) appear to offer metabolic enhancement, while higher doses (> 7mg/kg) produce the opposite effects and may even slow or inhibit mitochondrial respiration ^30^. However, the effects of MB on CNS may also be linked to its dual effect on oxidation (both anti- and enhancing), as well as the engagement with numerous other pharmacological targets. For example, MB is thought to be an inhibitor of soluble guanylate cyclase (sGC) and of nitric oxide synthase (NOS) enzymes ^31^, both of which modulate the NO-mediated vasodilation and could affect cerebral perfusion ^32^. Furthermore, MB is also a potent inhibitor of monoamine oxidase A (MAO-A) which could partially explain its proposed mood stabilising and anti-depressant properties ^33^. These and other molecular targets of MB have been reviewed elsewhere ^11^.

Despite the large body of literature supporting MB’s neurotropic effects in the CNS, data on how it affects the living brain are limited. Studies in rodents suggest that a low dose of MB (0.5 mg/kg) increases regional cerebral blood flow (CBF), oxygen extraction fraction (OEF) and cerebral metabolic rate of oxygen (CMRO_2_) under resting conditions ^34^ and even more so under hypoxic conditions coupled with sensory stimulation ^35^. These data provided initial corroboration of MB’s mitochondrial enhancing effects in animals *in vivo*, while the effects of MB on human brain oxygenation and metabolism have not yet been established. Interestingly, in contrast to the animal studies, Rodriguez et al. ^36^ reported a trend toward an (4 mg/kg) MB-induced *reduction* in human CBF, despite the apparent increase in network connectivity, providing contradictory evidence of *reduced* oxygen delivery to the tissue.

Given the paucity of animal data and a lack of direct evidence from human studies, we used quantitative neuroimaging to study the interactions of MB with CBF and metabolism *in vivo*, in both humans and rats. We reasoned that although preclinical evidence suggested that MB should enhance brain oxygenation and metabolism, owing to the known complexity of MB pharmacology, multiple mechanisms of action and suggested hormetic effects ^37^, as well as potential species-specific effects, a direct investigation and comparison of MB in both humans and animals was warranted.

We conducted two studies, in healthy humans and in rats, respectively. For the human study, we studied the effects of two doses of MB (0.5 mg/kg and 1.0 mg/kg) on quantitative MRI parameters. These included estimation of CBF and OEF, which enabled calculation of a parameter closely related to CMRO_2_. CMRO_2_ is a measure of the rate of the oxidative phosphorylation processes, and therefore represents an index of brain mitochondrial respiration efficacy. For the rat studies, we measured dose-dependent changes in CBF induced by 2 and 4 mg/kg of MB, during both stimulated and rest conditions. The rats were imaged under anaesthesia, as is customarily conducted in rodents to minimise stress, movement and regulate physiology. Indeed, others studying the effects of MB on brain of rats *in vivo* also used anaesthesia ^34, 35^. Instead of MR-based measurement of oxygen or glucose metabolism, which are technically more challenging at high magnetic fields (e.g. 9.4T used here) ^38^ we measured brain glucose metabolism by a classic method of ^14^C-2-deoxyglucose autoradiography (2DG) estimating the rate of glucose utilisation, CMRglu ^39, 40^. As we performed 2-DG in both awake and anaesthetized rats, this additionally allowed us to rule out the confounding interference of anaesthesia on our metabolic measurements.

Our main hypothesis was that MB would increase brain oxygen utilisation, by virtue of increase in CBF or OEF, or both. We also predicted a corresponding increase in CMRglu in rats. Such direct assessments of the neurometabolic effects of MB *in vivo* were expected to provide essential information to facilitate the therapeutic use of MB in neurological and psychiatric diseases underpinned by neuroenergetic deficits. Contrary to our expectations, we detected reductions in blood flow and metabolism in both species, highlighting the complexity of MB and the need for further investigations, which we also discuss below.

## Methods

### Human study

#### Study design and procedures

The study adopted a single-blinded, within-subject randomised design (Figure 1). Eight healthy volunteers received infusions of placebo (50 mL of 5% v/v glucose solution), and two MB doses (0.5 mg/kg and 1 mg/kg, each diluted in 50 mL of 5% v/v glucose solution), in three sessions, on separate days in randomised order. The infusions took place approximately 30 minutes prior to the MRI. Further details on participants selection and experimental methods are reported in the Supplementary Methods.

**Figure 1.**
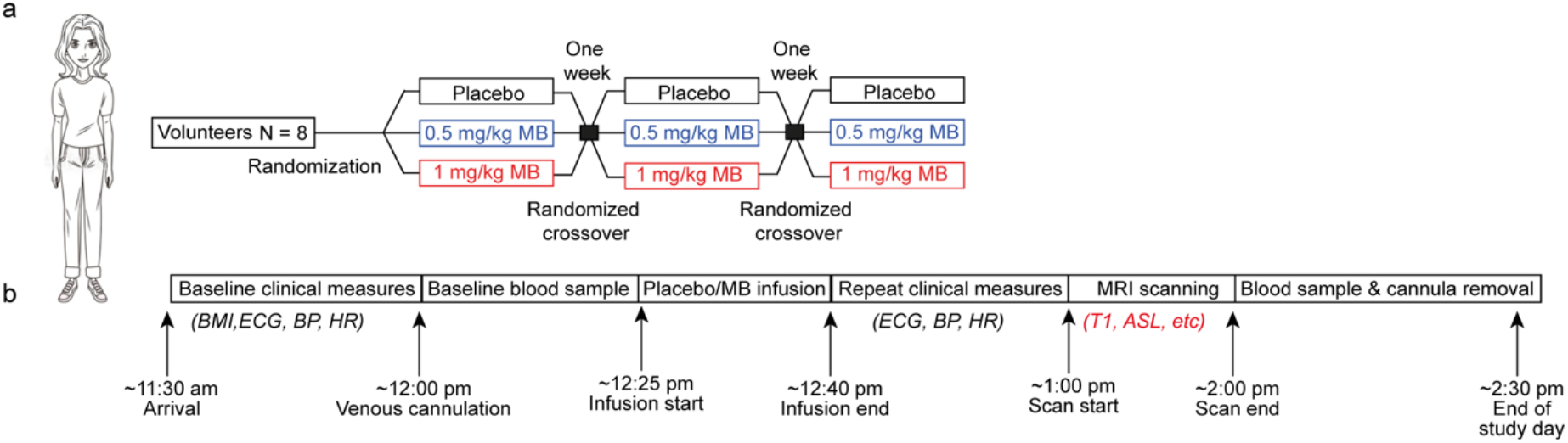
Illustration of the design of the human study.

Blood samples were collected on all three visits before and after MB infusion, and haematological parameters including red blood cells (RBC), mean corpuscular haemoglobin concentration (MCHC), packed cell volume (PCV) were measured. Additionally, blood pressure, electrocardiogram (ECG) and heart rate were also monitored pre- and post-infusions (results shown in Supplementary Table 1). Subjective ratings of Energy and Mood were obtained through the administration of Visual Analogue Scales, ranging from 0 to 100, after each infusion. Participants were asked to report any side-effects, including pain ratings on a scale of 0-10 (10 being the highest) experienced during the MB infusion. Subjective ratings are shown in Supplementary Figure 1.

#### MRI methods and analysis

MR images were acquired on a 3T GE Healthcare MR750 scanner (GE Medical Systems, Milwaukee, WI). The MRI protocol included high-resolution structural T_1_-weighted (T1w) magnetisation-prepared rapid gradient echo (MPRAGE) images, as well as measurement of regional cerebral blood flow (CBF) by 3D pseudo-continuous Arterial Spin Labelling (3D pCASL), and voxelwise R2’ and OEF by a quantitative BOLD method based on an Asymmetric Spin Echo (ASE) acquisition ^41, 42^. These techniques allow the quasi-simultaneous estimation of CBF and OEF respectively, which in turn enables the computation of a parameter closely related to CMRO_2_ that is proportional to their product. CBF maps were generated according to standard methods including those recommended by the ASL consensus paper ^43^. CMRO_2_ calculations were carried out using the method outlined by Yablonsky *et al*. ^44^, which is based on the Fick’s principle ^45^. Further details of MRI acquisitions and image processing are provided in the Supplementary Material.

### Rat study

#### Study design and procedures

The rat study consisted of *in vivo* MRI and autoradiography experiments. Both experiments adopted a between-subject randomised design. Animal experiments were in accordance with the UK Home Office Animals (Scientific Procedures) Act (1986) and were approved by the King’s College London ethical review committee.

To assess the effect of MB on rat brain perfusion and the BOLD response by MRI under anaesthesia, male adult Sprague-Dawley rats (Charles River, UK) were initially cannulated (tail and femoral vein, for MB and anaesthetic administration) under isoflurane anaesthesia, then switched to α-chloralose anaesthesia (65 mg/kg initial bolus, 30 mg/kg/hr infusion) for the duration of MRI scanning. This anaesthetic regime was chosen due to its well-documented property of preserving the neurovascular coupling, and hence BOLD response, during sensory stimulation paradigms ^46^. Each animal’s physiology was maintained and monitored throughout the experiment.

To assess the effect of MB on the rat brain metabolism under both anaesthetised and awake conditions, we undertook *in vivo* ^14^C-2-DG autoradiography in two cohorts of male adult Sprague-Dawley rats: one conscious and one anaesthetised with low-level isoflurane. ^14^C-2-DG enables a quantification of the regional brain glucose utilisation, in a matter analogous to ^18^F-FDG PET. The method involves timed administration of ^14^C-labelled glucose analogue deoxyglucose and simultaneous blood sampling. The radioactive brain tissue is subsequently sectioned and exposed to x-ray film for densitometric quantification. For full details of 2-DG procedure see Supplementary materials.

#### MRI methods and analysis

BOLD contrast sensitive images were acquired using a GE-EPI sequence, while CBF imaging used a continuous arterial spin-labelling (CASL) method. During the entire scanning protocol, non-noxious sensory forepaw stimulation and rest were alternated, providing BOLD images and CBF data from both resting and stimulated states. The entire combined BOLD/ASL protocol was repeated 5 minutes after intravenous administration of 2 (n=3) or 4 (n=4) mg/kg MB. Because we originally aimed to replicate the forepaw stimulation result on BOLD MRI ^35^ using 0.5 mg/kg MB, we additionally acquired BOLD (but not CBF) data with this dose (n=6). For full details of MRI protocols, sequence parameters for BOLD and ASL, sensory stimulation and drug/animal combinations see Supplementary materials.

#### *In vivo* autoradiography methods and analysis

All rats were initially anaesthetised with isoflurane to cannulate their femoral vein and artery. After this, animals undergoing conscious 2-DG were loosely restrained by wrapping their torso and lower limbs in plaster cast, and allowed to recover for 90 minutes to eliminate residual effects of isoflurane ^47^. Rats were intravenously administered 2 mg/kg MB, or vehicle, 30 minutes before ^14^C-2-DG to maximise the likelihood of MB accumulation in the brain ^12^. Brain glucose utilisation (GU) quantification was carried out according to the methodology previously described ^38, 40^ using MCID software (Interfocus, UK). Six ellipsoid ROIs across an entire hemisphere at 4 brain levels were averaged to estimate whole brain glucose utilisation.

## Results

### MB reduces CBF, OEF and CMRO_2_ in humans

Eight female, healthy participants took part in the study. They were non-smokers, with an average age 27 ± 8 years and had a body mass index of 22.5 ± 1.3 (mean ± SD). Haematological and other clinical parameters (blood pressure and ECG, and subjective ratings of Energy, Mood and Pain) are summarised in Supplementary Table 1 and Supplementary Figure 1.

We measured global CBF, OEF and CMRO_2_ after two doses of MB and after placebo (Figure 2). We observed significant MB-induced reductions in global CBF (F_(1.74, 12.17)_ = 5.82, p = 0.02). The 0.5 and 1 mg/kg doses of MB reduced CBF by 8.36% (p = 0.014), and 8.3% (p = 0.034) respectively, and no dose-dependent relationship was observed (p = 0.16). We also observed significant MB-induced reductions in global (grey matter) OEF in humans relative to placebo (F_(1.64, 10.67)_ = 6.761, p = 0.016). One value in the 0.5 mg/kg MB group was detected to be an outlier and was removed from this analysis. The 0.5 mg/kg MB dose did not significantly change OEF (−1.6% change relative to placebo, p = 0.17), whilst the 1 mg/kg MB dose caused a 2.7 % reduction in OEF (p = 0.032). OEF reduction by MB was dose-dependent (p = 0.0042).

**Figure 2:**
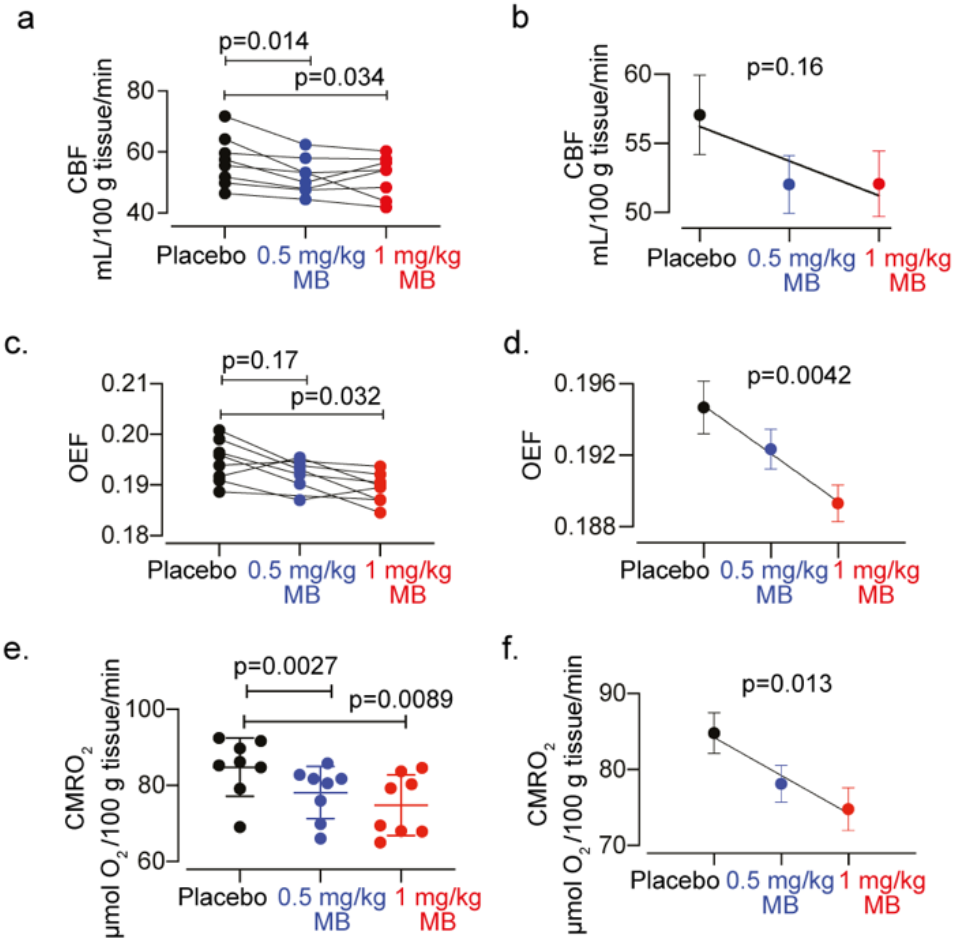
Effect of MB on CBF, OEF and CMRO2 in healthy human subjects (n=8). (a, c) Effects of 0.5 mg/kg and 1 mg/kg MB on CBF and OEF respectively, shown for individual subjects. (b) group CBF means ± SEM values and regression line. Both MB doses significantly decreased CBF but no dose-dependent effect was observed. (e) CMRO_2_ values derived from CBF and OEF for individual participants, with both MB doses showing significant reduction from placebo (d, f) group OEF and CMRO2 means ± SEM values and regression line. MB-induced reductions relative to placebo were dose-dependent.

MB treatment also significantly reduced CMRO_2_ relative to placebo (F_(1.26, 8.84)_ = 8.01, p = 0.016). The reductions in CMRO_2_ were 7.9% (p = 0.0027) and 11.8% (p = 0.0089) after 0.5 and 1 mg/kg MB, respectively. The CMRO_2_ reduction observed was dose-dependent (p = 0.013).

MB did not have any significant effects on blood pressure or heart rate, nor any of the haematological parameters, including the haematocrit, red blood cells or mean corpuscular haemoglobin concentration (MCHC) which are used in the calculation of CMRO_2_ (Supplementary Table 1).

### MB reduces CBF and glucose utilisation in rats

We measured resting CBF in anaesthetised rats (Figure 3), and we also measured cerebral glucose metabolism in both anaesthetised and conscious rats (Figure 4).

**Figure 3.**
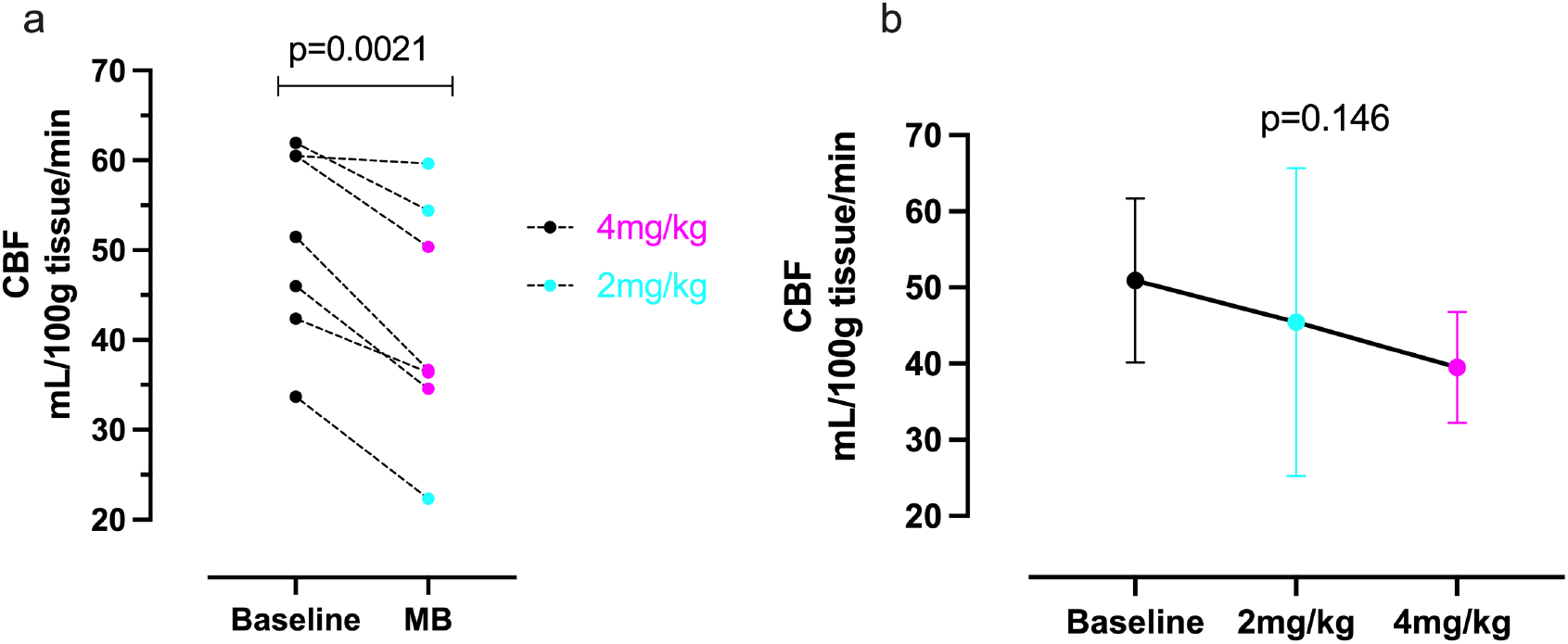
Effect of MB on CBF in rats (n=6). (a) Effects of 2 mg/kg and 4 mg/kg MB on CBF, comparing baseline and post-MB, in individual rats. (b) Group CBF means ± SEM values and regression line. Both MB doses significantly decreased CBF (paired t-test, p=0.0021) with a non-significant trend towards a dose-dependent effect (p=0.146)

**Figure 4.**
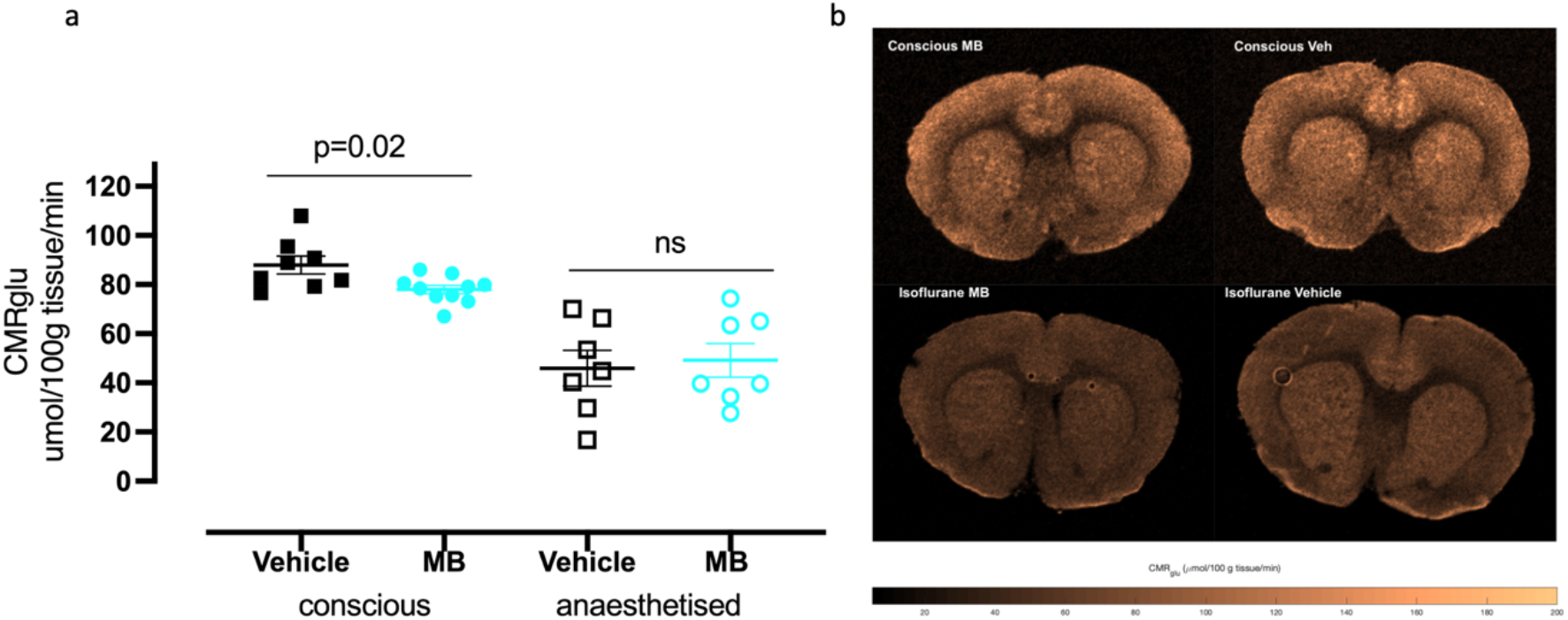
Effect of 2 mg/kg MB on global CMRglu in conscious and in anaesthetised rats. (a) There was a significant reduction of CMGglu in conscious rats after MB (p=0.02) but not in anaesthetised rats. (b) Quantitative CMRglu maps from one slice through the brain of four representative rats.

Because of the small sample size for each of the two groups in our CBF measurements, we combined both doses into one group and observed a significant reduction in global CBF in rats after MB, compared to before MB (paired t test, p=0.0021). The CBF reduction was 12.6% and 21.2% for 2 and 4 mg/kg, respectively (Figure 3a). Moreover, there was a trend towards a dose-dependent reduction in CBF (p = 0.146, 3b).

Using *in vivo* 2-DG autoradiography we next measured the global CMRglu, in response to 2mg/kg MB or vehicle, in both conscious and in anaesthetised rats. In the conscious rats, we observed a significant reduction in CMRglu with 2 mg/kg MB treatment (87.94 ± 10.26 μmol/100 g tissue/min) compared to vehicle (77.96 ± 5.54 μmol/100 g tissue/min; p = 0.018; Figure 4). We saw no significant difference between vehicle and MB in the anaesthetised rats (49.19 ± 18.05 MB, 45.96 ± 19.19 vehicle). It should however be noted that there was an overall ca. 50% decrease in the global CMRglu between conscious and anaesthetised rats, which is an expected effect of anaesthesia ^48^. In this case, the anaesthetised condition also increased the variance of the responses (coefficient of variation in vehicle rats increased from 9.4% in the conscious, to 39% in the anaesthetised), hereby decreasing the power to reject the null hypothesis and likely precluding the possibility to detect any effect of MB in this group of animals.

Representative maps illustrating the effect of MB on CBF and glucose utilisation in rates are also shown in Figure 4b. In summary, we observed a reduction in all brain CBF and metabolic parameters at rest in both humans and rats after a single administration of iv MB.

### MB does not increase stimulated CBF or BOLD response in the anaesthetised rats

To ascertain whether MB augmented neurovascular responsiveness to sensory stimulation, we measured both CBF and BOLD responses to non-noxious sensory forepaw stimulation, before and after MB. Forepaw stimulation is expected to induce increases in both CBF and in BOLD responses in the S1 forelimb cortex contralateral to the stimulated paw ^35^.

As expected, we observed a significant, 6.95% increase in contralateral (stimulated) S1 CBF (p=0.0045, Figure 5c) before MB. When repeating the stimulations after MB, the increases were smaller, 2.8% and 3% after 2 or 4 mg/kg dose of MB, respectively; when doses were combined, the overall increase in CBF of 2.94% was not significantly different from baseline (Figure 5c). There were no significant changes between stimulated and unstimulated CBF in the ipsilateral (unstimulated) S1, nor in the global (whole slice) CBF, irrespective of MB treatment, as expected.

**Figure 5.**
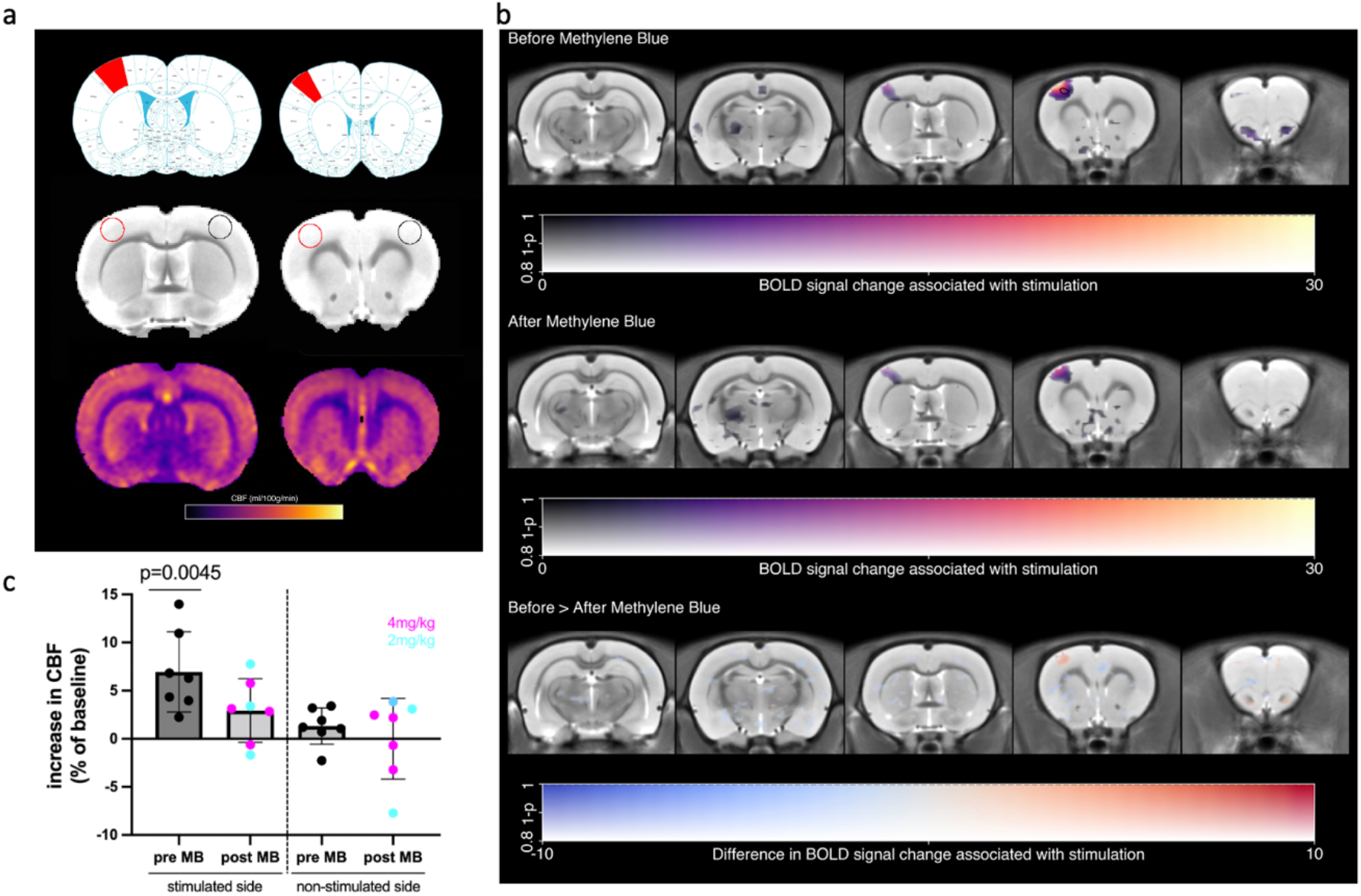
rat CBF and BOLD responses to forepaw stimulation. (a) ROIs (top - rat brain atlas showing location of forelimb S1 cortex (red); middle - structural images showing location of ROIs - red stimulated side, black non-stimulated side; bottom, representative CBF maps; (b) BOLD responses voxelwise for before (top) and after (middle) MB. The difference between the two conditions was tested (bottom) with a two-way ANOVA with MB dose and condition as factors. The response to stimulation is significant before MB (black contour denotes p<0.001 uncorrected), but not after, although the difference between the two conditions is not significant; (c) CBF % change from baseline, which was only significantly increased before, but not after MB, in the contralateral forelimb cortex (p=0.0045, one sample t test, compared to 0).

Raw CBF values from all treatment combinations are given in Table 1. In keeping with the aforementioned observations of decreased resting CBF (Figure 3), MB caused significant decreases in CBF in both S1 areas and in global CBF (post-vs pre-MB differences, Table 1).

**Table 1.** Resting (off) and stimulated (on) rat CBF (ml/100 g/min). #### p<0.0001 compared to pre; *p<0.05, ***p<0.001 compared to off. 2-way RM ANOVA, mean ± sd, n=7 per group (2 and 4 mg/kg MB groups combined)

|  | pre MB |  | post MB |  |
| --- | --- | --- | --- | --- |
| Region | *off* | *on* | *off* | *on* |
| **S1 ipsi** | 64±11.6 | 65±12.6 | 50.1±16.3 #### | 50.5±17.8 #### |
| **S1 contra** | 64±10.4 | 69±12 *** | 51.4±12.4 #### | 53.2±13.9 * #### |
| **Global** | 50.9±10.8 | 50.7±10.7 | 42.1±13.2 #### | 42.7±13.0 #### |

To characterise the effect of MB on BOLD responses resulting from forepaw stimulation, we also performed voxelwise, whole brain statistical parametric mapping analysis (Figure 5b). We observed a characteristic pattern of significantly increased BOLD response in the contralateral S1 area in stimulated rats before MB administration (Figure 5b; top panel); this cluster appeared after MB administration but was no longer statistically significant (Figure 5b; middle panel). However, the difference between the MB conditions was not significant (Figure 5b; bottom panel).

Taken together, the results from rats indicate that MB caused a decrease in both resting CBF and in glucose metabolism, in direct support of our CBF and CMRO_2_ data from humans. Moreover, and contrary to expectations, MB also did not increase stimulation evoked CBF or BOLD responses, robustly ruling out the originally proposed enhancement of neurovascular coupling in animals.

## Discussion

We report the results of quantitative imaging experiments designed to assess the effects of MB on CBF and metabolism in healthy humans and in rats. To the best of our knowledge, this is the first study to report quantifiable changes in human cerebral metabolism in response to MB, despite its clinical use since the 17^th^ century.

Contrary to our original expectations, we show robust reductions of resting CBF after MB administration in both species. We detected a ~8% decrease in global human CBF, after 0.5 – 1 mg/kg MB (Figure 2), and 12 and 20% decrease in global rat CBF after 2 and 4 mg/kg MB, respectively (Figure 3). All these results are in sharp contrast with the findings previously reported by Lin et al. ^34^ who demonstrated an 18% global CBF increase in anaesthetised rats 10 minutes after 0.5 mg/kg MB iv administration. To the best of our knowledge, other than the aforementioned Rodriguez *et al*. ^36^ who demonstrated reduced stimulation induced CBF in humans, no other human studies have examined acute effects of MB.

We further characterised brain metabolism in both humans and in rats, and we show MB-induced decreases in both. Specifically, we demonstrate that both doses of MB decreased CMRO_2_ in humans, and that the higher (1 mg/kg) dose also significantly decreased OEF; all parameters also show a significant dose dependency (Figure 2). In conscious rats, 2 mg/kg of MB significantly decreased CMRglu, which is a gold-standard direct measurement of brain metabolism. We did not see the same decrease in anaesthetised rats, but we note that anaesthetic-induced decrease in metabolic rate and a consequent decrease in SNR might have reduced our power to detect an effect. These results are, again in contrast to those by Lin *et al*. ^34^ who demonstrated significant increase in both CMRglu and CMRO_2_ in rats after 0.5 mg/kg MB, albeit both in anaesthetised animals. Finally, and contrary to Huang at al. ^35^ we did not detect increased BOLD response to somatosensory forepaw stimulation after neither MB dose they originally used (0.5mg/kg; Supplementary Figure 2) nor after our 2 and 4 mg/kg doses (Figure 5). In other words, in our study MB (0.5-4mg/kg) did not appear to potentiate stimulated BOLD response in the rat brain.

Having based our *a priori* hypothesis on MB stimulating mitochondrial oxygen consumption and ATP production as suggested by both *in vitro* and animal studies ^29, 34^, the observed reduction in all cerebral metabolic measures, in both humans and rats, was unexpected. However, various aspects of MB’s pharmacology provide crucial hints for the interpretation of our findings and may explain the discrepancies from the results reported by others. MB’s pharmacological action is biphasic: within the hormetic zone, MB enhances mitochondrial-dependent energy production and ATP synthesis and *in vitro* evidence from rat brain homogenates show MB doses between 0.5 and 5 μM increase cytochrome C activity; however doses higher than 10 μM result in cytochrome C activity inhibition, possibly due to oxidising effect of MB on mitochondrial electron transport chain ^37^.

Clinical observations confirm these hormetic mechanisms and are also relevant to *in vivo* observations. Use of MB as treatment of methaemoglobinemia indicates antagonistic effects depending on the concentration; i.e, doses up to 4 mg/kg iv convert methaemoglobin to haemoglobin, but doses exceeding 7 mg/kg, which are considered cytotoxic appear to worsen methemoglobinemia ^11, 49^. Behavioural pharmacological studies in rodents report that intermediate MB doses (between 5-20 mg/kg) are associated with maximal behavioural response consistent with memory enhancement effect, while does below 5mg/kg and over 20 mg/kg have either none or opposite effects ^37^. It is worth noting the species differences between humans and rodents where, due to smaller surface to volume ratio and higher metabolic activity, estimated equivalent dose in rats to humans is approximately 6x higher ^50^; hence we chose 2 & 4 mg/kg to approximately match our (0.5 and 1 mg/kg) human doses while still in keeping within dose range previously shown to affect brain metabolism in rats.

MB’s poly-pharmacological actions include mechanisms other than its cycling reducing-oxidising action. Of particular relevance to our study, MB can inhibit nitric oxide-induced vasodilation via blockade of guanylyl cyclase activation and NOS antagonism ^11^. Nitric oxide-induced vasodilation is one of regulatory mechanism of CBF ^51^, which enables the brain to adjust to increased oxidative phosphorylation and metabolic demand by increasing supply of substrates, namely oxygen and glucose, in response to increased neuronal and glial energetic requirements. If MB antagonises NO-induced vasodilation this could alter normal neurovascular responses. Again, dose consideration is important as MB has little effect on the nitrergic system at very low doses, whilst its anti-vasodilating effect increases with the dose ^31^.

Based on these arguments, it is plausible to predict that MB concentrations at the lower end of the hormetic zone may cause an increase in CMRO_2_ resulting from stimulation of oxidative phosphorylation processes, with resulting coupled haemodynamic effects, i.e. an increase in CBF and CMRglu. This would be consistent with the findings of Lin *et al*. ^34^ study which used a very low (0.5 mg/kg) dose in rats. However, once MB concentrations raise nearer or beyond the limit of the hormetic zone range, MB’s cumulative effect could be determined by a progressively lower stimulation of the oxidative phosphorylation metabolic rate, and by larger NOS inhibition, resulting in a null or even inhibitory effect on CBF and metabolism.

It is therefore possible that in our human and animal study, the concentration of MB in the brain was different to those previously used in aforementioned studies and may be too high for the expected metabolic enhancing effect. In support of this theory is the fact that a trend toward *decrease* in fMRI task-related CBF in humans was observed by Rodriguez *at al*. ^36^ using a higher dose of 4 mg/kg. The doses we used are in keeping with the current clinical practice, and most beneficial therapeutic effects MB in treatment of methemoglobinemia, cyanide poisoning, and malaria, do correspond to MB doses of 1 to 2 mg/kg.

Despite this, the difference in concentrations between blood and CNS needs to be taken into account. Peter *et al*. ^12^ demonstrated an accumulation of MB in CNS resulting in MB concentrations 20 times higher in the brain relative to blood. Considering they administered 1 mg/kg iv resulting in an approximate plasma concentration of about 2 μM, it might be extrapolated that our dose of 0.5 - 1 mg/kg in humans is very likely to cause a much higher MB concentration in the brain (plausibly in a range between 10 to 40 μM) relative to the concentrations in blood known to induce antioxidant and pro-metabolic effects. Nevertheless, we also measured BOLD responses in rats using 0.5 mg/kg and we did not detect increases after MB, in direct contrast to Lin et al. ^34^ and Huang at al. ^35^ who used the same dose in rats.

Interestingly, and in support of our hypothesis, we noticed that the lower MB dose used in the human study was associated with higher energy ratings relative to placebo and the higher dose. There was no effect of any MB dose on mood relative to placebo. Pain ratings instead increased linearly with the dose. We did not observe significant changes in blood related parameters, i.e., the haematocrit, red blood cells or mean corpuscular haemoglobin concentration (MCHC) (Supplementary Table 1). Whilst we could not monitor O_2_ saturation in our human study, as MB interferes with the light absorption dependent SatO_2_ signal measured using pulse oximetry, we were able to monitor pO_2_ throughout scanning in rats by pulse oximetry and the values were within normal range between 97-99%.

There are some limitations in our study. We have only recruited female subjects in our human study, whereas the rats were males, both of which limits the generalisability of our findings, although we are not aware of any evidence of sex-specific effects of MB. Furthermore, our sample size is very small, although the consistent findings we observed across subjects indicates that the direction of the effects observed in our study is not the result of an underpowered analysis.

The OEF values obtained by ASE, and consequently the CMRO_2_ values derived from the product of CBF and ASE, are lower than those detected in the human brain using older and more validated methods. Discrepancies between ASE-derived OEF estimates and those obtained with invasive and ^15^O_2_ PET methods, are likely deriving to the overestimation of deoxygenated blood volume (DBV) known to result from the implementations of the qBOLD model we used ^52^. However, this should have not affected our findings due to the study’s within-subject, cross-over design. We have analysed OEF at whole-brain level and therefore we cannot provide analysis of regional distribution of OEF, and consequently of CMRO_2_, changes. OEF maps obtained with ASE are characterised by low signal to noise (SNR) ratio^42^, and analysis of smaller regions would be problematic and reduce confidence in our results obtained in a small sample. Future studies in larger samples, and using improved modelling of qBOLD data to address OEF underestimation (e.g. by the inclusion of Z-shimming to account for macroscopic field inhomogeneities), and to obtain better SNR of OEF maps, will be needed to explore the distribution of the effects of MB in the brain.^53^ Moreover, based on current results we can only infer on acute MB administration effects. Our findings do not preclude the possibility that the chronic use of MB may lead to metabolic enhancements, in line with preliminary evidence of clinical efficacy in a 6 month trial in Bipolar Disorder ^17^.

In conclusion, we report robust human and rat imaging data suggestive of an inhibitory effect of MB on brain metabolic measures, at doses used clinically for treatment of haematological conditions. Data from clinical trials application of MB provide further support to our hypothesis that the most beneficial effects of MB at CNS level is expected using extremely small MB doses, significantly lower than used for treatment of methaemoglobinemia. Interestingly, a clinical trial of an MB derivative with putative anti-TAU aggregating properties (hydromethylthionine) demonstrated that the lower dose (8 mg/kg) originally meant to be used as control, had significant pro-cognitive and anti-brain atrophy effect, whilst the test doses of 200 mg/day were not associated with any benefit on those parameters ^54–56^. In an earlier study, investigating mood stabilising effects of MB, a 15 mg/day oral dose originally selected as placebo condition, equating to ~0.2 mg/kg, and 20 times lower than the putative active dose, was found to be potentially clinically effective ^57^.

In conclusion, MB, at the doses employed in clinical haematological applications, reduces cerebral perfusion and energy metabolism in rats and humans, at rest. It is possible that the proposed neurometabolic enhancement effect of MB requires lower doses, in both healthy volunteers and in rodents. Further quantitative imaging studies of MB effects on cerebral blood flow and metabolism are necessary to characterise its potential in therapeutic CNS applications.

## Acknowledgements

We are grateful to the radiographers of the Centre for Neuroimaging Sciences. We thank Mattia Baraldo for his helpful assistance with references, and to Tobias Wood for help with rat study MRI protocols.

## Author contributions

AC, FT, FZ, DC conceived the studies

AC, NS, FZ, DL, DC designed the studies

NS, AC, KR, NM, PS, SS carried out experimental procedures for the human study

OD, DL, FZ carried out imaging analyses for the human study

EM, CS, DC carried out experimental procedures and imaging analyses for the rat study

NS, AC, EM, DC and FZ wrote manuscript draft

All authors approved the submitted manuscript

## Disclosure / Conflict of interest

### Funding

The human study was partly funded by the Mental Health Biomedical Research Centre award of the King’s College London and the South London & Maudsley NHS Trust and by a NARSAD Young Investigator grant from the Brain and Behavior Research Foundation to AC. The rat study was funded by the UK Biotechnology and Biological Sciences Research Council (BB/N009088/1). EM was supported by the UK Medical Research Council (MR/N013700/1) and King’s College London as a member of the MRC Doctoral Training Partnership in Biomedical Sciences. FET was funded by the NIHR Maudsley Biomedical Research Centre at South London and Maudsley NHS Foundation Trust and King’s College London. There are no conflicts of interest.

## Notes

### Competing Interest Statement

The authors have declared no competing interest.

