## Supplemental materials for "The effects of acute Methylene Blue administration on cerebral blood flow and metabolism in humans and rats"

### Supplementary Methods

#### Human study

Participant information and ethics

Ethical approval was granted from the Psychiatry, Nursing and Midwifery Research Ethics Subcommittees, King's College, London (Ref: HR-16/17-4804) and was sponsored by King's College, London. Informed consent was obtained from all participants prior to conducting any study procedures.

Exclusion criteria included any current medical condition, current or previous psychiatric or neurological condition, glucose-6-phosphate dehydrogenase (G6PD) deficiency, pregnancy, taking any drug that may act via the serotonin system, and lack of MRI safety.

Methylthioninium chloride or MB (ProveBlue®, Martindale Pharmaceuticals Ltd.), injection ampules (5 mg/mL), were purchased from the hospital pharmacy. Methylene blue was diluted, as approprtiate, in 5% glucose before intravenous administration.

Placebo and Methylene Blue solutions were administered through a syringe pump intravenously via a cannula placed in the median cubital vein. The three sessions took part a week apart in randomised order, at approximately the same time of day (13:00 ± 1 hours). The participants were blinded to the dose of methylene blue administered. The rate of infusion was 4.1 mL/min ± 0.1 (mean ± SD) and was identical for each infusion for each participant. The duration of the infusions ranged between 12 and 15 minutes (check). MRI acquisition began ~ 33 minutes after the start of the infusion (32.7 ± 9.7 mins, mean ± SD).

MRI protocol and analysis

MRI images were acquired using a 3T GE Healthcare MR750 scanner (GE Medical Systems, Milwaukee, WI).

Radiofrequency was transmitted with the scanner body coil, while signal was received with a 12-channel receive-only head coil.

After an initial localizer scan, high-resolution anatomical images were acquired using an adapted 3D T1-weighted magnetization-prepared rapid gradient echo (MPRAGE) sequence with the following parameters: 1.2-mm isotropic resolution, repetition time of 7.312 ms, echo time of 3.01 ms, and inversion time of 450 ms.

CBF data were acquired using 3D pCASL to determine changes in regional resting perfusion. The sequence included background suppression for optimum reduction of the static tissue signal, which increased sensitivity to the labelled arterial blood signal. The labelling RF pulse had a duration of 1.8s and a post-labelling delay of 2.025s. Four control-label pairs were collected, each within ~45 s, to derive a mean perfusion weighted difference image. After the postlabeling delay, images were acquired using a multishot, segmented 3D fast spin echo stack-of-spiral sequence (8-arms) with an effective spatial resolution of 2 × 2 × 3 mm. A proton density (PD) image was also acquired using the same acquisition parameters to enable the computation of quantitative CBF maps [1].

Asymmetric Spin Echo (ASE) data were acquired with TR/TE= 3100/74ms, 10 repetitions, t=4ms, 16 echo-shifts. Only the first 8 shifts were considered optimal for the R2' and OEF estimation.

CBF maps were produced from ASL and PD images according to methods recommended by the ASL consensus paper [2]. Raw PD and T1w images were co-registered, and transformations were applied to CBF maps which were also normalised to MNI space using SPM12 software [3]. Normalised CBF maps were smoothed using an 8-mm FWHM kernel. Global CBF values in the grey matter were extracted with the Functional Software Library suite (FSL; [4]) and were compared across conditions using non-parametric tests (Wilcoxon signed ranks test). Voxel-wise comparisons of the CBF maps across treatments was performed using permutation tests, with and without global CBF covariance.

Voxel-wise R2' and OEF values were estimated from the ASE data, according to Blockley and Stone [5, 6]. Field maps were also acquired to correct for macroscopic magnetic field inhomogeneities [5].

Computation of $CMRO_2$

$CMRO_2$ calculations were carried out, using the method outlined by [7], which is based on the Fick's principle[8]. Briefly, $CMRO_2$ was calculated via the equations below:

*Equation 1: CMRO2 = CRBC CBF Hct Ya OEF*

Where $C_{RBC}$ is the oxygen carrying capacity of blood and is a product of the mean corpuscular haemoglobin concentration (MCHC) and the amount of oxygen that can be carried by 1 g of pure haemoglobin, which is a known practical quantity of 1.34 mL/g. $Y_a$ is the oxygenation level of arterial blood, which we assumed to be the reasonable value of 0.98 for healthy subjects. Therefore, we adapted Equation (1) to calculate CMRO2 for each participant as shown below in Equation (2).

*Equation 2: CMRO2 = CBF OEF Hct 1.34 MCHC 0.98*

Finally, this was simplified as

*Equation 3: CMRO2 = CBF OEF Hct MCHC 1.31*

Hct was measured along with MCHC and other blood parameters on each visit, as well as before and after each treatment. CBF and OEF were obtained from the ASL and ASE scans, as outlined above. Our MRI-derived CBF units were mL blood/100g tissue/min, therefore the values we obtained for $CMRO_2$ had units of mL $O_2$/100g tissue/min. This was converted to the more universally used $CMRO_2$ units which are μmol $O_2$/100 g tissue/min by converting mL $O_2$ to μmol $O_2$, as 1 mol of gas corresponds to 22.4 L.

#### Rat study

##### Animal information and ethics

All animal experiments were conducted in accordance with the Home Office Animals Scientific Procedures Act, UK, 1986 (Project licence P023CC39A) and were approved by the King's College London ethical review committee. Adult (10-16 weeks old), male Sprague–Dawley rats, Charles River, UK) were group-housed at 21 ± 1 °C in a 12-h light:dark cycle and with ad libitum access to standard rat chow and drinking water. Animals were housed for a minimum of one week prior to any experimental procedures. Animals were assigned to either dose of MB, or to vehicle treatment, using simple randomisation. The reporting of this study complies with the Animal Research Reporting *in vivo* experiments (ARRIVE) guidelines [9].

##### MRI protocol and analysis

Rats were initially anaesthetised with 2.5-3% isoflurane in an 80:20 mix of air to medical oxygen for cannulation of the tail and femoral veins. Isoflurane was discontinued (while air and oxygen continued to be administered) and anaesthesia switched to α-chloralose by injecting a bolus of 65 mg/kg via the tail vein cannula before infusion at a rate of 30 mg/kg/hr. Body temperature, respiration rate and pulse oximetry (SA Instruments, Inc; Stony Brook, NY) were monitored throughout the experiment.

Data were acquired using a 9.4 T Bruker Biospec MR scanner with 86 mm volume, four channel array receiver, and arterial spin labelling coil. The scanning protocol consisted of a structural scan, followed by 10 minute BOLD protocol, 12 minutes CBF protocol, iv MB administration and 5 minutes later, repetition of BOLD and CBF protocols. BOLD response was measured using a GE-EPI sequence with the following parameters: TR 1000 ms, TE 18 ms, $\alpha$ 90°, 24 slices, 64 x 64 matrix with voxels 0.35 x 0.35 x 0.8 mm). Cerebral blood flow (CBF) measurements were derived from the pairs of tagged and control images using the continuous arterial spin-labelling (CASL) method with the following parameters: TR 4000 ms, TE 20 ms, $\alpha$ 90°, 96 x 96 matrix, single 1 mm slice, 8 seconds per volume.

During the scanning we also applied somatosensory, non-noxious electrical stimulation to the right forepaw via a plantar TENS pad and dorsal platinum needle with the following stimulation parameters: 200mV, 3 Hz, 0.4 ms, 2 mA [10]. We alternated periods of rest and stimulation as follows: for BOLD scans, 10 epochs of 40 seconds with no stimulation and 20 seconds with stimulation, resulting in total of 600 brain volumes (400 resting, 200 stimulated), and for CBF scans it was ten epochs of 48 seconds with no stimulation and 24 seconds with stimulation, resulting in single slice CBF maps with 60 volumes at rest and 30 stimulated (sCBF) volumes. This 22 minute protocol was repeated five minutes after administration of either 2 mg/kg (n=3) or 4 mg/kg (n=4) iv MB (1 mL/kg). Additionally, six rats underwent BOLD protocol only, and were administered 0.5 mg/kg MB.

The structural image was corrected for inhomogeneous signal intensity using N4BiasFieldCorrection [11], and normalised to a rat template image [12] with antsRegistration [13]. BOLD volumes were aligned to the first volume with AFNI's 3dVolReg [14, 15]. The mean image across time was calculated from the motion-corrected volume which was subsequently registered to the structural image. Finally, the motion-corrected volume was warped to the template image by concatenating the transformations from BOLD-to-structural and structural-to-template. A general linear model was fit to the boxcar model from the forepaw stimulation paradigm convolved with a haemodynamic response function and the realignment parameters from motion correction for each subject. Random effects were analysed with a full factorial analysis of dose (2 or 4 mg/kg) and MB status (before or after MB). Contrasts positively associated with stimulation were generated with one-sided t-tests which were statistically assessed at the group level with a two-tailed factorial design and the pTFCE toolbox [16] for SPM12 [3].

CASL single slice images were corrected for motion artefacts by registering the timeseries to the mean volume, before absolute CBF images were calculated using qi-asl from Quantitative Imaging Tools [17]. Mean CBF time courses were extracted from the whole slice (whilst avoiding its edges), and two bilateral ellipsoid ROIs in the somatosensory cortex using Jim8 software (Xinapse Systems, UK).

*In vivo* autoradiography protocol and analysis

Rats in the autoradiography experiment (n=32) were anaesthetised with isoflurane in order to cannulate the femoral vein and artery. Animals for the conscious 2DG cohort (n=18) were then restrained by wrapping their torso and lower limbs with a loose-fitting plaster cast (3M), and once this had hardened, the anaesthesia was terminated. The rats were monitored for absence of stress visually and by measuring their plasma glucose periodically, ensuring it is within normal physiological values. Animals from anaesthetised 2DG cohort remained under a low dose of isoflurane (1-1.2% in air/oxygen 80/20). At 90 minutes from discontinuing anaesthesia (in conscious animals), or ca. 120 minutes after starting the anaesthesia, all rats were intravenously administered 2 mg/kg MB or vehicle. After 30 minutes, $^{14}$C-2-DG (Perkin Elmer, USA) was infused over 30 seconds intravenously and 14 timed arterial blood samples collected as previously described (Littlewood et al., 2006). After the final blood sample at 45 min post-$^{14}$C-2-DG, the animals were decapitated and the brains rapidly removed, frozen in cold isopentane (~-40°C) and stored at -80°C. Quantification of plasma glucose and $^{14}$C was carried out using the blood glucose analyser (YSI 2300) and scintillation counter (Beckman Coulter LS 6500), respectively. Brains were cryosectioned at 20 µm and exposed to x ray film (Kodak Biomax MR-2) alongside calibrated $^{14}$C standards (American Radiolabelled Chemicals) for 7 days, after which they were developed in Optimax 2010 x-ray film processor. Brain glucose utilisation (GU) quantification was carried out by calculating µmol/100g/min from the optical densities in films, and from plasma glucose and

$^{14}C$, according to the methodology originally described by Sokoloff *et al.* [18] using MCID software (Interfocus, UK). Six large ellipsoid ROIs across an entire section at approximately 1, -3, -6 and -8 mm from Bregma [19] were averaged to estimate whole brain glucose utilisation.

#### Statistics

All statistical analyses (human and rat experiments) were performed with Graphpad Prism 8 (version 8.1.1) and all reported p-values have been adjusted for multiple comparisons using the Holm-Sidak test for multiple comparisons. All data was tested for normality using the D'Agostino & Pearson test, outliers were tested using the Grubbs method ($\alpha$=0.05).

For the human data, determination of statistical significance was carried out using a repeated measured one-way analysis of variance (ANOVA) with a Geisser-Greenhouse correction, or in case of incomplete data sets, due to outliers, a mixed analysis model was used.

For the rat CBF data (Figure 3), 2 and 4 mg/kg MB doses rats were combined and a paired t test used to ascertain the differences between pre and post-MB scans.

Also, one sample t test was used to demonstrate significance of percent changes from baseline (Figure 5a). For each rat, CBF data was collected from both the left and right hemispheres and then averaged. For the conscious autoradiography (Figure 4), Mann Whitney t-tests were used to determine statistical significance; those were corrected for multiple comparisons using Holm Sidak method.

Additionally, to compute dose-related trends, a simple linear regression analysis for trend was also carried out to determine any significant dose-dependent effects for the relevant human and animal data.

Unless otherwise mentioned, all error bars in the figures are standard error of mean (SEM). The data shown in Table 1 are mean ± standard deviation (SD).

All image analyses were conducted by the operators blinded to the identity of the group or drug dose.

### Supplementary Tables and Figures

**Supplementary Table 1. The clinical parameters measured showed no significant treatment-related differences. The values shown in the table are mean ± SD, N=8.**

| Clinical measure | Placebo (5% glucose) | Low dose (0.5 mg/kg MB) | High dose (1 mg/kg MB) | p value RM one-way ANOVA |
|---|---|---|---|---|
| **Blood** | | | | |
| Hb (g/L) | 131.3 ± 12.15 | 132.5 ± 13.33 | 130.4 ± 13.41 | 0.70 |
| Hct (L/L) | 0.41± 0.035 | 0.42 ± 0.039 | 0.41 ± 0.041 | 0.73 |
| MCHC (g/L) | 318.9 ± 5.55 | 318.4 ± 6.42 | 319.2 ± 4.75 | 0.89 |
| RBC (L/l 10^12) | 4.42 ± 0.39 | 4.44 ± 0.38 | 4.36 ± 0.21 | 0.56 |
| **Blood pressure** | | | | |
| Systolic BP (mm Hg) | 113.1 ± 8.34 | 111.9 ± 8.02 | 114.9 ± 8.46 | 0.34 |
| Diastolic BP (mm Hg) | 73.31 ± 10.49 | 74.00 ± 5.13 | 76.00± 8.52 | 0.36 |
| Heart rate (bpm) | 65.94 ± 8.24 | 61.31 ± 9.86 | 64.50± 12.52 | 0.45 |
| **ECG** | | | | |
| QTC | 412.7 ± 27.08 | 404.6 ± 14.49 | 410.9 ± 17.78 | 0.96 |
| PR | 150.3 ± 24.94 | 151.3 ± 20.31 | 156.5 ± 20.36 | 0.17 |
| QT | 395.1 ± 26.43 | 404.3 ± 33.13 | 402.3 ± 37.74 | 0.23 |
| QRS | 79.43 ± 5.74 | 81.50 ± 7.98 | 79.75 ± 7.29 | 0.25 |

**Supplementary Table 2. Number of rats per group and mean weight ± SD per experimental group**

| Group | 2 mg/kg | 4 mg/kg | 0.5 mg/kg | vehicle | Mean weight (g) |
|---|---|---|---|---|---|
| **MRI** | | | | | |
| BOLD and CBF (stimulated and resting) | 3 | 4 | | | 467.1 ± 136.7 |
| BOLD only (stimulated) | | | 6 | | 342.0 ± 28.2 |
| **2DG** | | | | | |
| anaesthetised | 7 | | | 7 | 321.0 ± 28.0 |
| conscious | 10 | | | 8 | 336.4 ± 16.4 |

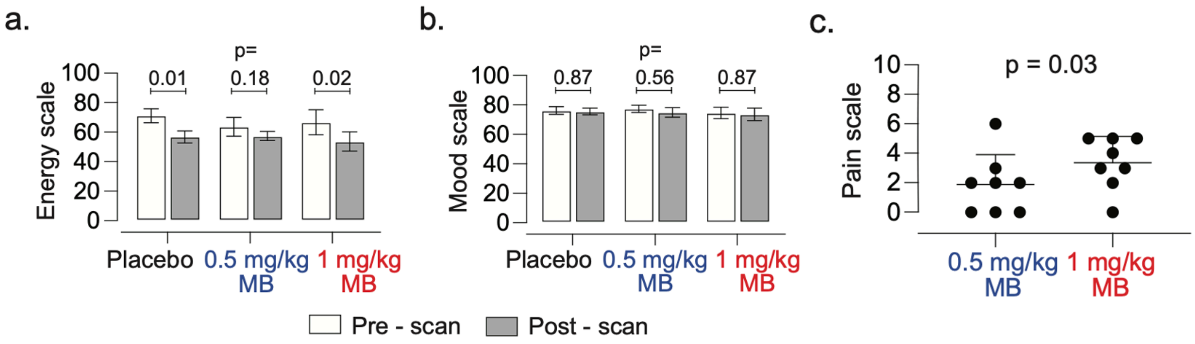


**Supplementary Figure 1: subjective effects**

(a, b) Bar graph showing visual analogue scale ratings for all participants before and after the infusion of placebo, 0.5 mg/kg and 1 mg/kg methylene blue. There was a statistically significant reduction in energy ratings after the placebo and 1 mg/kg scans, but not after the 0.5 mg/kg scan. No difference was observed in mood before and after the scans for any of the drug administrations. (c) Scatter plot showing the pain ratings at the time of infusion for the 0.5 mg/kg and 1 mg/kg methylene blue. There was a statistically significant increase in pain from the lower to the higher dose. No pain was reported during the infusion of placebo.

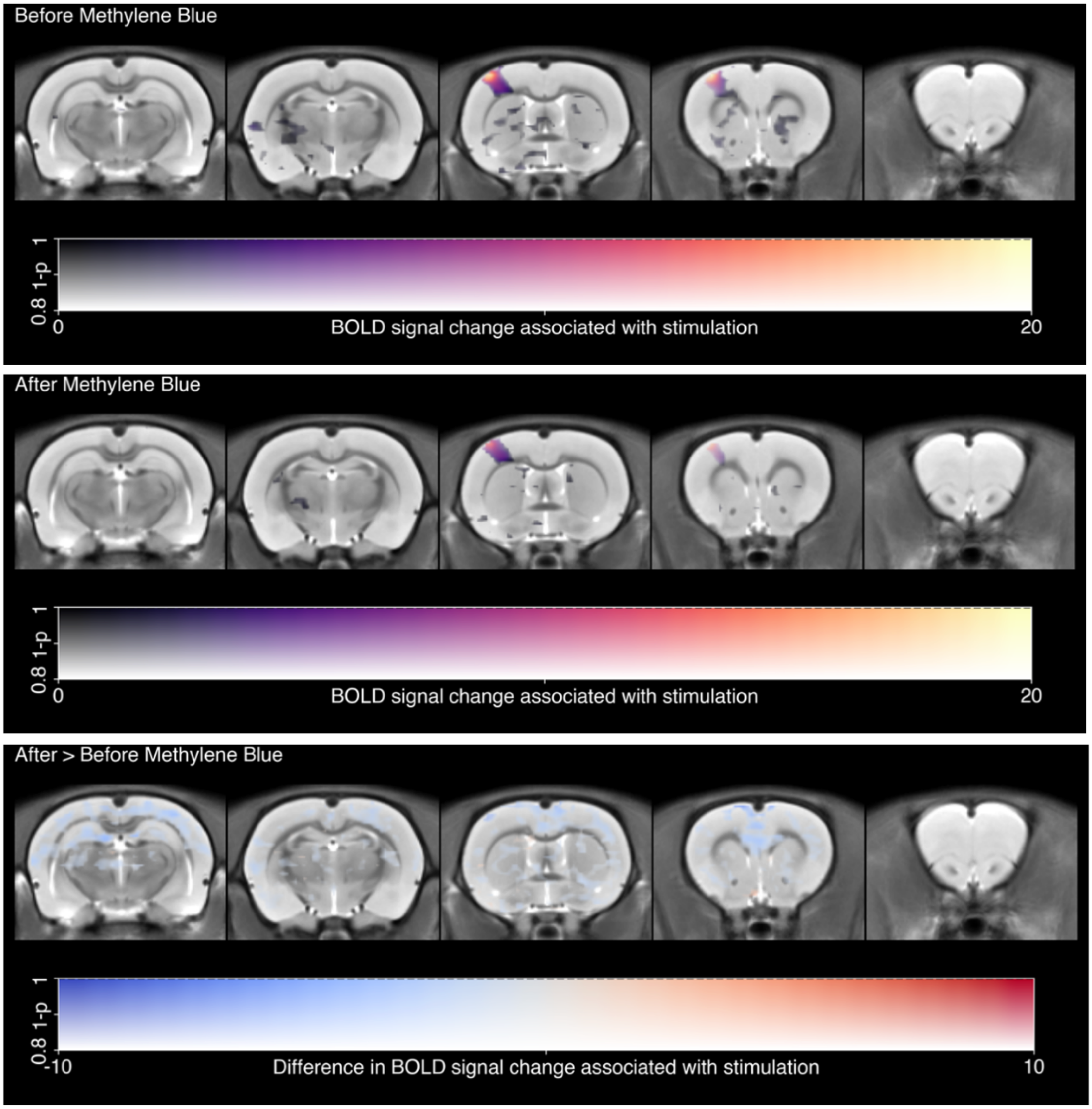


**Supplementary Figure 2: Voxelwise BOLD responses to non-noxious electrical stimulation of the forepaw before (top) and after (middle) 0.5 mg/kg of MB.**

The difference between the two conditions was tested with a paired two-tailed T-test (bottom). The response to stimulation did not surpass the significance criteria of $p<0.001$ at the second (group) level, but significant somatosensory clusters were present in the first (individual) level (data not shown). Nonetheless, there was no significant differences between before and after MB.
